# Mutualistic interaction with *Trichoderma longibrachiatum* UENF-F476 boosted plant growth-promotion of *Serratia marcescens* UENF-22GI

**DOI:** 10.1101/2020.08.24.265587

**Authors:** Régis Josué de Andrade Reis, Alice Ferreira Alves, Pedro Henrique Dias dos Santos, Kamilla Pereira Aguiar, Silvaldo Felipe da Silveira, Luciano Pasqualoto Canellas, Fábio Lopes Olivares

## Abstract

**BACKGROUND:** A plethora of bacteria-fungal interactions occurs on the extended fungal hyphae network in soil. The mycosphere of saprophytic fungi can serve as a bacterial niche boosting their survival, dispersion, and activity. Such ecological concepts can be converted to bioproducts for sustainable agriculture. Accordingly, we tested the hypothesis that the well-characterized beneficial bacterium *Serratia marcescens* UENF-22GI can enhance their plant growth-promoting properties by combination with *Trichoderma longibrachiatum* UENF-F476.

**RESULTS:** The colony and cell interactions demonstrated *S. marcescens and T. longibrachiatum* compatibility. Bacteria cells were able to attach, forming aggregates-biofilms and migrates through fungal hyphae network. Bacterial migration through growing hyphae was confirmed using two-compartment Petri dishes assay. Fungal inoculation increased the bacteria survival rates into the vermicompost substrate over the experimental time. Also, *in vitro* indolic compound, phosphorus, and zinc solubilization bacteria activities increased in the presence of the fungus. In line with the ecophysiological bacteria fitness, tomato and papaya plantlet growth was boosted by bacteria-fungi combination applied under plant nursery conditions.

**CONCLUSION:** Mutualistic interaction between mycosphere-colonizing bacterium *S. marcescens* UENF-22GI and the saprotrophic fungi *T. longibrachiatum* UENF-F467 increased the ecological fitness of the bacteria alongside with beneficial potential for plant growth. A proper combination and delivery of compatible beneficial bacteria-fungus represent an open avenue for biological enrichment of plant substrates technologies in agricultural systems.

## INTRODUCTION

Plant growth-promoting bacteria (PGPB) have been used as a technological alternative to reduce the use of fertilizers and pesticides in agriculture (1, 2). The positive effects of the bioinoculants application in agriculture are partly dependent on the ability of the microbial agent to colonize the rhizosphere and the host plant under the influence of the physical-chemical parameters of the soil-plant system. Also, the soil microbial community is highly diverse and competitive (3), limiting the establishment of bioinoculants formulated with strains selected under laboratory and greenhouse conditions (4, 5).

Different technological approaches have been used in order to formulate more efficient inoculants, which must be combined with suitable application methods in the agricultural environment (1, 6–8). An adequate formulation must guarantee a minimum number of viable microbial cells, and once applied in the soil-plant system, establish itself in the rhizosphere or plant-host body. Thus, the inoculant formulation must guarantee the survival of the cells for a period that allows their shelf-life commercialization (6). Also, the rapid decline of the introduced microbial population, reduce competitiveness, and plant colonization capability (9, 10). Among the technological proposals aimed to increase the survival and activity of a specific microorganism introduced into the soil, we can mention the use of polymeric carriers, stabilized organic matter and its sub-fractions, organic additives, encapsulation, and combination with other microorganisms (1, 2, 11, 12).

Basic and applied aspects of interactions between endomycorrhizal or ectomycorrhizal fungi with bacteria are explored in the literature (13–15). Less attention is given to interactions involving saprotrophic fungi and bacterial communities, despite their pivotal roles in biogeochemical cycles, nutrient cycling, and organic matter dynamics (16–18).

Saprotrophic fungi are eukaryotic microorganisms endowed with survival structures (spores), which germinate and colonize the environment when conditions are favorable and with high capacity of environment dispersion, migrating through hyphae tip-growth to different niches. Ecological studies have shown the benefits of fungi on the survival, dispersion, and activity of bacterial communities in the soil, rhizosphere, and plant tissue (5, 19–21).

Different bacteria species can colonize the micro-habitat called sapro-rhizosphere (12) that encompassing the mycosphere and mycoplane of fungi hyphae, and which serve as a biotic surface for physical anchoring, chemical signal exchange and nutritional niche (12, 20–23). Mycosphere competent-colonizer bacteria do not depend on the oligotrophic C-soil sources or highly competitive C-exuded by the plants in the rhizosphere (20). Other competitive advantages are the maintenance and dispersion of bacteria in the environment (22, 24, 25). Furthermore, soil fungi can degrade complex substrates (26) and mutually benefit the ecological interaction by metabolic cooperation and successional communities in the diverse microbial process.

Overall ecological advantages suggest that certain saprotrophic fungi may be selected to enhance the ecological fitness and beneficial properties of PGPB that give rise to several potential technological applications on bacterial-fungal interactions. In the present study, a well-characterized plant growth-promoting bacterium *Serratia marcescens* (Sm-22GI) strain UENF-22GI (27) and the saprotrophic fungi *Trichoderma longibrachiatum* (Tl-F476) UENF-F476 were combined to test potential enhancement of the bacteria ecological fitness and plant growth-promoting properties *in-vitro* and *ex-vitro*. The taxonomic position of the fungi at species level was defined herein, and the compatible colony-colony physical interaction between Sm-22GI and Tl-F476 was previously demonstrated (28). Both microorganisms were isolated from matured cattle manure vermicompost, being subject to test in combination to prove the technical feasibility of the biological enrichment of plant substrates (biofortification concept).

## MATERIALS AND METHODS

### Microorganisms used and inoculum production

Both microorganisms were isolated from cattle manure at 90-days (fungal isolate) and 120-days (bacteria isolate) mature vermicompost using earthworms (*Eisenia foetida*). Genome sequencing and plant growth-promotion traits of *Serratia marcescens* strain UENF-22GI were detailed reported at Matteoli and colleagues (27).

The purified bacterium was stored in glycerol 10% at −80°C and activated by inoculation of 100 μL of the bacterium stock in glass tubes containing 5 mL of Nutrient Broth (NB) liquid medium (8g L^−1^) and incubated in a rotatory shaker at 30°C and 150 rpm for 48-h. The Sm-22GI inoculum was prepared by centrifugation and resuspension of the cell in sterilized water at OD_(492 ηm)_ = 1.0 with 2 × 10^8^ cells mL^−1^. This procedure was repeated twice. The Tl-F476 fungal isolate was stored in a glass tube containing sterilized vermiculite and activated in the potato-dextrose liquid medium after incubated in a rotatory shaker at 28 °C and 150 rpm for 3-days. After that, 100 μL of the suspension was transferred for potato-dextrose-agar (BDA) solid medium, followed by incubation at 28°C for 7-days, where the agar plug was obtained, or spore were resuspended in sterilized water and adjusted on Neubauer chamber for 5 × 10^6^ spore mL^−1^.

### Molecular taxonomic identification of the fungus isolate F476

The saprotrophic fungus isolate F476 was previously classified as *Trichoderma* sp. based on the colony and cell morphology, respectively, on PDA solid medium and differential and interferential contrasting light microscopy (DIC).

For molecular identification of fungi, genomic DNA was extracted from the pure colonies using the hyphae tip method (29), and the *internal transcribed spacer* (ITS) region was used since it has a high phylogenetic signal for identification of *Trichoderma* sp. Amplification reactions were performed with the primers specific for ITS1 and ITS4 (30, 31). The purified samples were sent for sequencing on the company ACTGene Análises Moleculares (Biotechnology Center, Federal University of Rio Grande do Sul, Porto Alegre, RS, Brazil). The identification of the fungus isolate UENF-F476 followed the protocol suggested by the International Commission on Taxonomy of *Trichoderma and Hypocrea* (ISTH), using the program TrichOKEY (32). The isolate was allocated to the *Longibrachiatum* section, and a phylogenetic study was carried out within this section to confirm the identification (Supplementary Table 1). Phylogenetic reconstruction was created using Bayesian Inference (BI) employing the Markov Monte Carlo chain method (MCMC) was performed. MrMODELTEST (33) was used to select the nucleotide replacement model for BI analysis. The likelihood values were calculated, and the model was selected according to Akaike Information Criterion (AIC). The evolution model selected for ITS was HKY + I + G. The BI analysis was completed with MrBayes v.3.1.1 (34), using 10^7^ generations. The trees were sampled every 1,000 generations, resulting in 10,000 trees. The first 2,500 trees were discarded from the analysis. The following probability values (35) were determined from the consensus tree through the remaining 7,500 trees. The convergence of the likelihood logs was analyzed with the software TRACER v. 1.4.1 (36). The species *Penicillium glabrum* SQU-QU09 was used as an external group (outgroup) in the analyzes.

### Evaluation of *in vitro* colony and cellular compatibility *S. marcescens* and *T. longibrachiatum*

The physical colony-colony compatibility between Sm-22GI and Tl-F476 was demonstrated. Curiously, the same strain was effective in countering two phytopathogenic fungi (27, 28). The previous results were confirmed by evaluating the same strains over ten days (10-d) incubation time, according (27). We also tested the phytopathogen *Fusarium oxysporum* against Sm-22GI.

The cellular interaction was carried out according to (37) with few modifications. Briefly, one 5 mm diameter agar plug containing fungal mycelium was placed on the center of a glass slide that was previously autoclaved at 121°C for 15 minutes. Four aligned spots containing 2 μL of the bacterial suspension (~ 10^6^ cells per spot) were inoculated on the glass slides at 2 cm equidistant points at each side of the fungus disc. The assay was carried out inside sterile Petri dishes lined with moistened autoclaved filter paper and incubated for seven days at 28°C. Daily, the hyphae growth was monitored using an inverted light microscopy Zeiss Axio Vert.A1 and the physical interaction between fungal hyphae and the bacterium photo-documented using Zeiss Axioplan light microscope equipped with an AxioCam MRC5digital camera by interferential and phase-contrast techniques.

### Evaluation of *in vitro* migration of *S. marcescens* via growing hyphae of *T. longibrachiatum*

The bacterium ability to migrate carried by fungal hyphae growth was accessed according to Rudnick and colleagues (38) with few modifications. In a bipartite Petri dish, a 5 mm diameter agar plug of the fungus mycelium was placed on the left side Petri dish compartment contained 2% agar-water solid medium poor-nutrient (PM). Five aligned and 1 cm equidistant spots of 10 μL of the bacterial suspension (~ 2 × 10^6^ cells per spot) were placed at 2 cm distant from the fungi agar plug. Nutrient Broth solid rich medium (RM) was added to the right compartment. The plates were incubated in a growth chamber for 7-d at 28° C. Daily evaluation was done to evaluate if the hyphae grew, crossed the plate barrier, and carried the bacteria cells for the right-side compartment (RM). At 3, 5, and 7 days after inoculation, agar plugs (10 mm diameter) were taken from the rich medium compartment to evaluate the bacteria co-migration. The samples were gently agitated in 2 mL of saline solution. The microbial suspension was filtered in a five μm-pore size (MF-Millipore™), and the retained material was washed with a 9 mL sterile saline solution. The whole volume suspension (10^−1^) was subjected to serial tenfold dilution followed by plating on NB solid medium containing 50 μg mL^−1^ of cycloheximide for bacteria count using four replicates per harvest time. The plates were incubated in the growth chamber for 4-d at 30° C, and CFU was determined. Data were submitted to log _10_ transformations. Control for inoculation of only bacteria or fungal on poor nutrient compartment was performed with no *S. marcescens* detection on the rich medium compartment after 7-d assay.

### *S. marcescens* survival on mature vermicompost in the presence of *T. longibrachiatum*

The survival of Sm-22GI bacterium was accessed in mature cattle manure vermicompost in the presence and absence of the isolated Tl-F476 over the 112-d period. The final inoculum density was adjusted for 5 × 10^6^ spores mL^−1^ and 5 × 10^8^ cells. mL^−1^, respectively for the fungi and bacteria and 100 μL of each suspension was applied in a sterilized glass vial (13 mL) containing 1 g of vermicompost (experimental unit), sealed with cotton plugs kept on the bench at room temperature. The experimental design was entirely randomized with three treatments: non-inoculated control, Sm-22GI, and Sm-22GI plus Tl F476, and three biological replicates per treatment were used, and samples were collected at 1 hour, 14-d, 28-d, 56-d, and 112-d after inoculation. The population size of the Sm-22GI bacteria was evaluated by diluted 1 g of the vermicompost in 9 mL sterile saline solution (NaCl 0.85%). After 15 minutes of agitation (50 rpm), 1 mL of the 10^−1^ dilution was transferred to a glass tube with a 9 mL saline solution to obtain 10^−2^ dilution and successively performed until 10^−8^ dilution. From all serial dilution (10^−2^ to 10^−8^), 100 μL suspension was taken and spread with Drigalski loop onto NB solid plates, using three replicates per dilution. The population density was estimated by colony-counts of typical *S. marcescens* and expressed as population size % of the initial population level (100%), where the survival decline can be easily noted. Each time harvest was pair-wide compared to evaluate the significance of the bacteria survival related to the *Trichoderma longibrachiatum* presence.

### *S. marcescens* production of derived auxins and phosphate solubilization capacity in the presence of *T. longibrachiatum*

Both microbial inoculums were produced as described above. Suspensions of 25 μL of the Sm-22GI, Tl F476, and their combinations were transferred to test tubes containing 5 mL of DYGS liquid medium (39) with and without the addition of tryptophan (100 μL of the 100 mg L^−1^ stock solution in distilled water and filtered through 0.22 μm Millipore membrane, then stored in the dark in a refrigerator). The test tubes were incubated in the dark for 72 h at 30 °C and 150 rpm under orbital shaker. Production of derived auxin (40) was estimated by transferring 150 μL of the supernatant to a 96-well polystyrene microplate, with 100 μL of Salkowski’s reagent (1 mL FeCl_3_.6H_2_O (0.5 mol) being added L^−1^) in 50 mL of HClO_4_ perchloric acid (35% in water), with an incubation period of 30 minutes in the dark. The reading was performed using a Hidex Chameleon Multilabel Detection Platform spectrophotometer with 492 nm absorbance by the MikroWin 2000 program. Increasing concentrations from 0, 6.25, 12.5, 25.0, 50.0, 100.0, and 200 μM of IAA diluted in the DYGS medium were used to construct the calibration curve. The concentration was measured using the calibration curve, relating absorbance and AIA concentration. Three repetitions were performed for each treatment and data expressed in μM IAA-equivalent.

For evaluation of P-solubilization, 100 μL suspensions od Sm-22GI, Tl F476, and their combinations inoculum were transferred to 50 mL test tubes containing Pikovskaya liquid medium (41), supplemented with 1 g L^−1^ of tribasic calcium phosphate (CP) [Ca_3_(PO_4_)_2_] or with 1 g L^−1^ of Araxá rock phosphate (RP). The test tubes remained under constant agitation on the orbital shaker at 150 rpm for 10-d at 28 ° C. Afterwards, they were centrifuged at 3200 rpm for 15 min, and the supernatant was used to quantify the levels of soluble phosphorus by the colorimetric method of ammonium molybdate, with absorbance reading at λ = 600 nm. The treatments control medium RP and CP, Sm-22GI, Tl F476, and their combinations with three biological replication per treatment.

### *S. marcescens* inoculation effect on tomato (*Solanum lycopersicum*) and papaya (*Carica papaya*) plantlets under greenhouse conditions in the presence of *T. longibrachiatum*

The Sm-22GI bacterium inoculum was obtained as previously described. The Tl F476 biomass was produced using rice as an inoculum vehicle, which was previously sterilized for 15 min at 121°C for three times with 24-h interval for each autoclaving. PDA solid medium BDA plaque previously grown with the isolate UENF-F476 was washed with 50 mL of sterile distilled water, thus obtaining spores and mycelia of the fungus in suspensions, being inoculated in 500 g of rice in polyethylene bags. The rice was incubated for 15 days at 28 °C. At 24 h intervals, the rice bag was shaken for homogeneous growth of the fungal isolate F476. In 1 g of rice, with approximately 1 × 10^6^ conidia g^−1^, 1 ml of the bacterial suspensions (10^8^ cells mL^−1^ was added under constant agitation, thus obtaining the mixed inoculants. Tomato (Santa Cruz, giant Kada) and papaya (Caliman-01 - UENF) seeds were sown together with inoculants in the substrate formulated with 50%v cattle manure vermicompost and 50%v washed sand washed in pots with a volume of 280 cm^3^. The controls received the same portion of uninoculated sterilized rice. The seedlings were harvested in 20 and 25 days after sowing for the tomato and papaya, respectively. The plant biomass was evaluated by measurement of the fresh and dry mass of the aerial part and the root (expressed mg plant^−1^). The experimental design was completely randomized with six treatments and seven replicates. The treatments were as follows: T1 = uninoculated control; T2 = *Trichoderma* F476; T3 = *S. marcescens* UENF-22Gi; T4 = Combined application of *S. marcescens* + *Trichoderma* F476.

### Statistical analysis

Statistical analyses of the data obtained were performed using one-way variance analysis (ANOVA at a confident level of 5%) that is conclusive for pair-wise comparison. The Tukey test compared data averages with more than two treatments at 95% confidence.

## RESULTS

### Identification of the *Trichoderma* sp. isolate UENF-F476

The fungal isolate coded UENF-F476, obtained from 90-d matured cattle manure vermicompost, was recognized as a member of the *Trichoderma* genus based on the colony and cell morphology (Fig. 1). Phylogenetic analysis was performed with 32 taxa, and the alignment of the sequences resulted in a total of 744 characters, of which 85 were informative for parsimony, 219 were variable, and 475 were conserved. Phylogenetic analysis using the ITS gene made it possible to identify the isolate in the *T. longibrachiatum* clade, well supported (pp = 0.96) (Fig. 2), confirming the previous identification by the TrichOKEY software, which allocated the isolate understudy in the same section. The overall analysis points to the strains UENF-F476 as a representative of the *Trichoderma longibrachiatum*.

**Fig 1.**
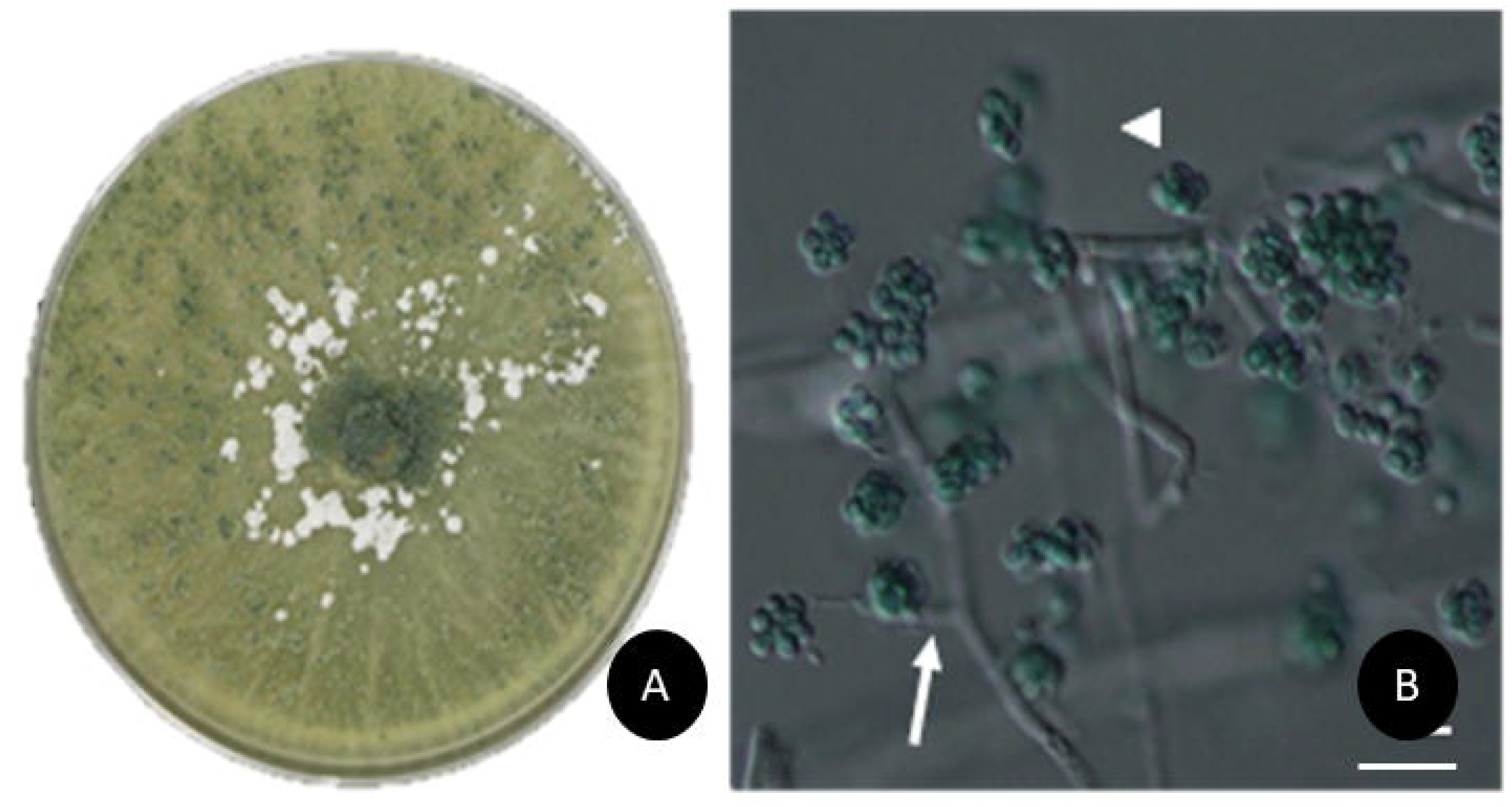
The colony of *Trichoderma longibrachiatum* strain UENF-F476 in potato dextrose agar (PDA) (**A**). Interferential microscopy of microculture technique sporulation (**B**) showing conidia (arrow-head) and conidiophores (arrow), bar = 20μm.

**Fig 2.**
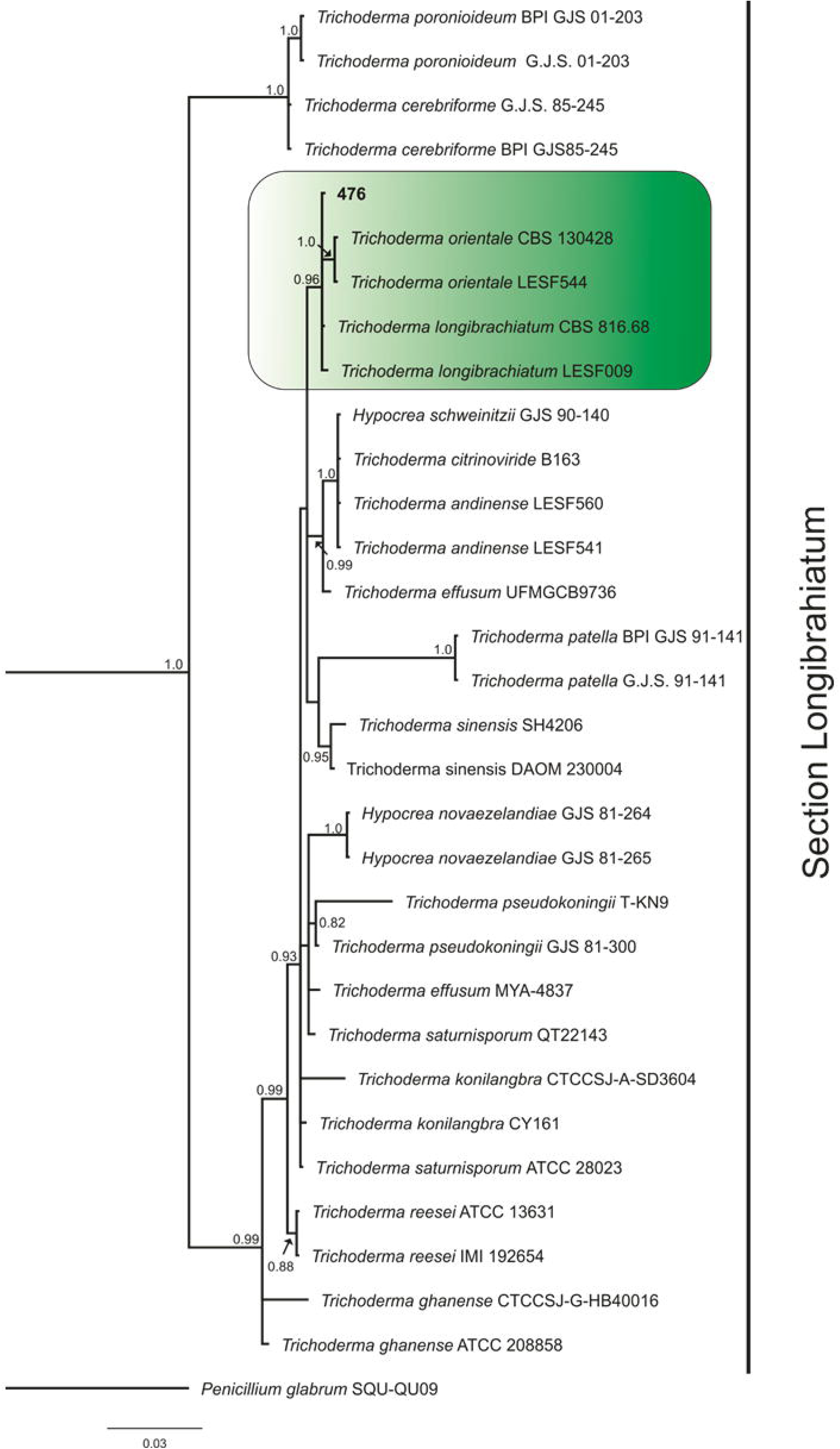
Phylogram based on Bayesian Inference of ITS gene sequences from isolates of *Trichoderma* sp. The posterior probability is indicated close to the branch nodes. The tree was rooted in *Penicillium glabrum* SQU-QU09.

### Colony-colony compatibility of the *S. marcescens* and *T. longibrachiatum*

The compatible physical contact between the colonies of Sm strain UENF-22GI and Tl strains UENF-F476 was previously demonstrated (27, 28). The assay was repeated with longer evaluation time and proper control (Fig. 3). After 2-d incubation, it was possible to visualize fungal hyphae growing in the direction of *S. marcescens* colonies (Fig.3-B1). After 6-d, fungal hyphae colonized the whole plate surface area intersected with bacteria colonies (Fig. 3-B2c). Both microorganisms were able to grow alone abundantly in the controls (Fig. 3-B2a,b) with a reduction of Sm colony-size, probably due to nutrient competition. Interestingly, when a mixture of Sm strain UENF-22GI and Tl strains UENF-F476 was spotted on the center of the plate, the hyphae kept growing stained red, due to *S. marcescens* pigmentation (Fig, 3-B3). A distinct phenotype was obtained when *F. oxysporum* fungal was challenged by Sm, resulting in pigmentation around the fungal colony with growth restriction (Fig. 3-A2, A3).

**Fig 3.**
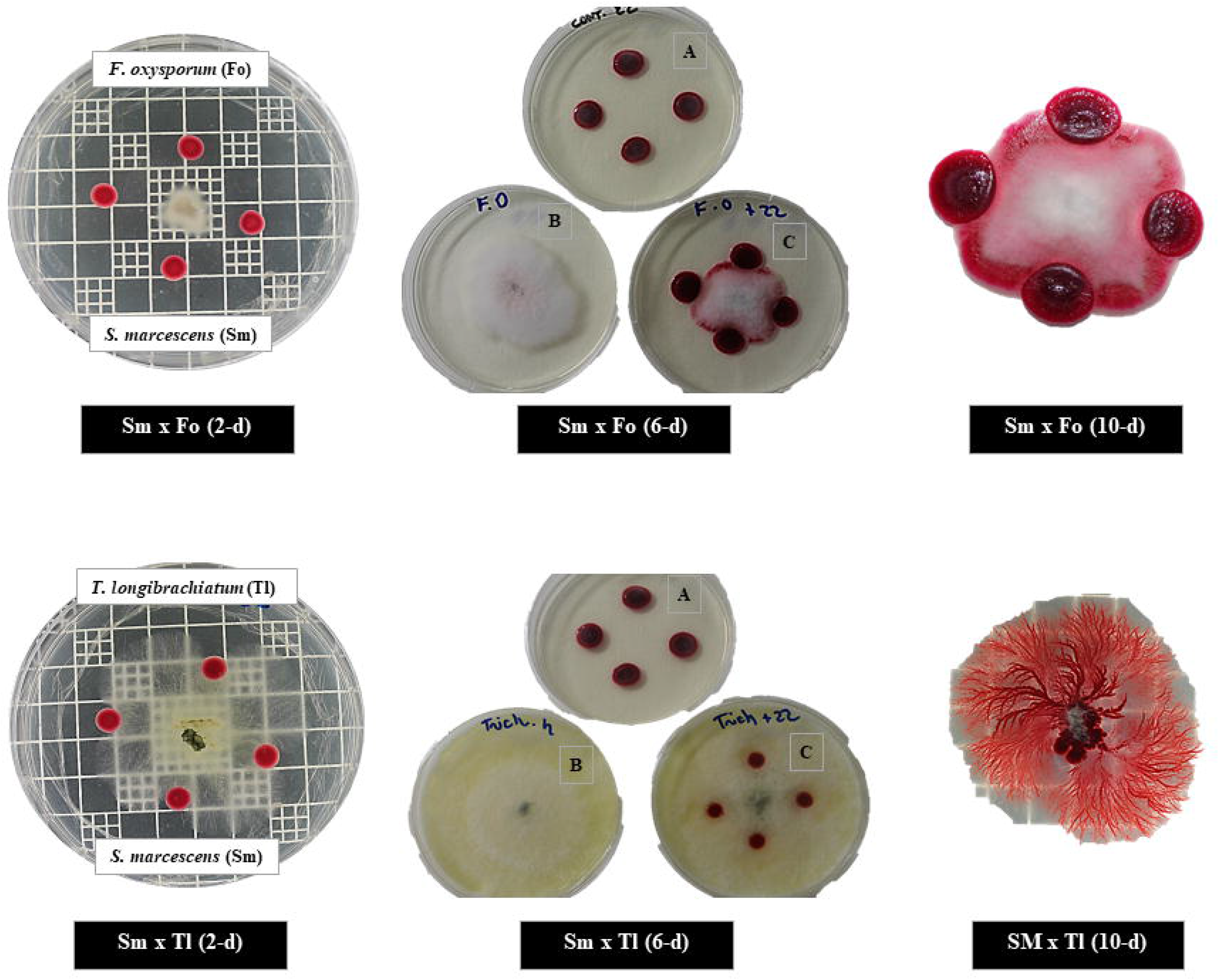
Colony compatibility of *S. marcescens* UENF-22GI and *T. longibrachiatum* UENF-F476 on NB solid medium. The growth pattern of the fungal hyphae showing physical compatibility with *S. marcescens* over 6-days assay (B1-B2). Note the *T. longibrachiatum* naturally stained by red due to the bacteria colonization and pigmentation of the growing hyphae (B3). When *F. oxysporum* interacted with *S. marcescens*, the fungal growth was limited by the bacteria presence (A1-A3).

### Cellular compatibility and interaction of the *S. marcescens* and *T. longibrachiatum*

It was possible to observe the interaction between *S. marcescens* strain UENF-22GI and *T. longibrachiatum* strain F476 (Fig. 4). The hyphae (h) growth was visible with 1-d after placing the agar plug on the glass slides, expanding in the direction of the bacteria (b) spot (Fig. 4A). Hyphae that emerged from bacteria colonies were permanently associated with the bacteria and continually explored the environment (Fig. 4B). Interesting to quote that hyphae became reddish pigmented, indicating the presence of the bacterium and released prodigiosin pigment (Fig. 4C). After seven days of interaction, the adhesion of bacteria on the cell-wall fungal surface was evident by forming small aggregates and migrating associated to the tip-growth hyphae (Fig. 4D-E). The main site for single bacteria cell attachment was the septal region of expanding hyphae (Fig. 4F). Thus, new hyphae exploring new environments remained associated with discrete aggregates of *S. marcescens* at regular intervals coinciding with the tip-growth and septa, which are probably regions of metabolite efflux (Fig. 4D-F). In regions with an older hyphae network, it was possible to observe the remarkable colonization of the hyphosphere by bacterial aggregates and biofilms, the increasing population size and survival on the environment (Fig. 4G).

**Fig 4.**
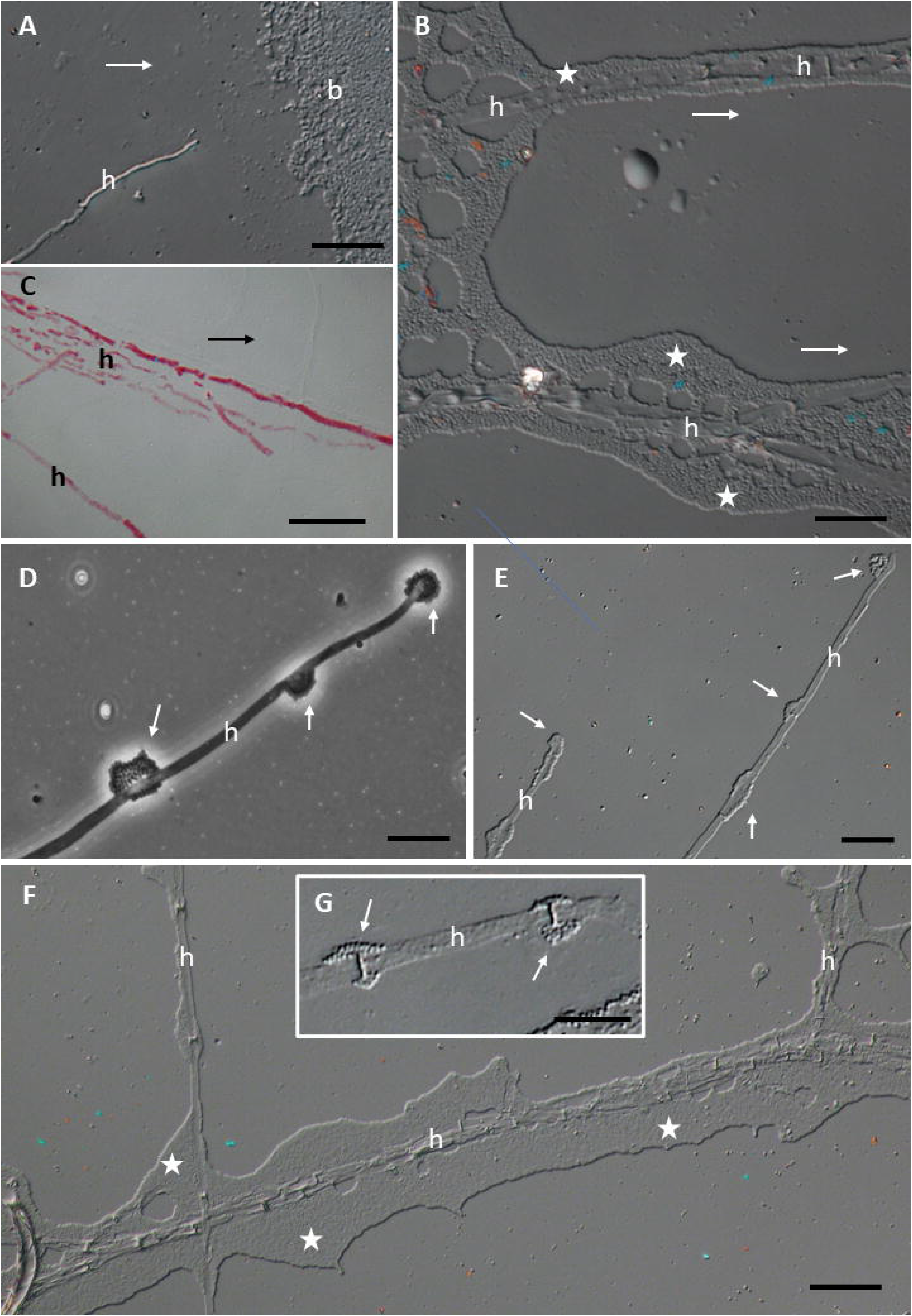
Differential interferential contrast (A, B, D-G), bright field (C), and phase-contrast (D) light microscopy of the interaction between *S. marcescens* UENF-22GI and *T. longibrachiatum* UENF-F476 on glass slides over a 7-d period. (A) Hyphae were approaching (h) bacteria spot (b) at 1-d after inoculation. (B) Hyphae (h) emerging from bacteria spots carrying bacteria aggregates alongside the hyphosphere and hyphoplane (white star). (C) Fungal hyphae are stained by reddish prodigiosin pigmentation released by the bacteria. (D-E) freshly growing hyphae were showing small bacteria aggregates migrating through the environment. Aggregates are associated with the tip-growth and laterally colonizing the hyphae by regular intervals (black arrows). (F) Discrete aggregates are attached to the hyphal septa region (white arrows). (G) Old hyphae (h) network massively colonized by bacteria aggregates and biofilms (white star). A-I Scale bar equal 50, 150, 100, 15, 50, 50, and 40 μm.

### Migration assay of the *S. marcescens* mediated by *T. longibrachiatum*

Using a bi-partite Petri dish, we evaluated the possibility of the growing fungal hyphae transport bacteria cells from a poor-nutrient to a rich-nutrient compartment. The hyphae growth was visible in 1-d, and after 3-d, the fungal crossed the barrier and colonized the rich medium (RM) compartment (Fig. 5). Prominent colonization and sporulation of the isolate F476 were observed in RM, as noted from 5-d to 7-d (Fig. 5). The fugal movement from PM to RM was accomplished by bacteria cell transport, as reported by the microscopical studies herein. The population size of *S. marcescens* recovered from RM-partition increased with the time from 3.7 to 6.8 log_10_ CFU. The results demonstrated that the fungal hyphae passed through bacteria spots and transported them from PM to the RM compartment. When the bacteria were inoculated in the poor medium without fungus (B-Control treatment), it was not possible to detect *S. marcescens* on the rich medium compartment remained after 7-d, demonstrating the role of the fungus as a bacteria carrier. The same result was obtained for fungal inoculation with bacteria (F-Control treatment), where no *S. marcescens* counts were obtained (Fig. 5).

**Fig 5.**
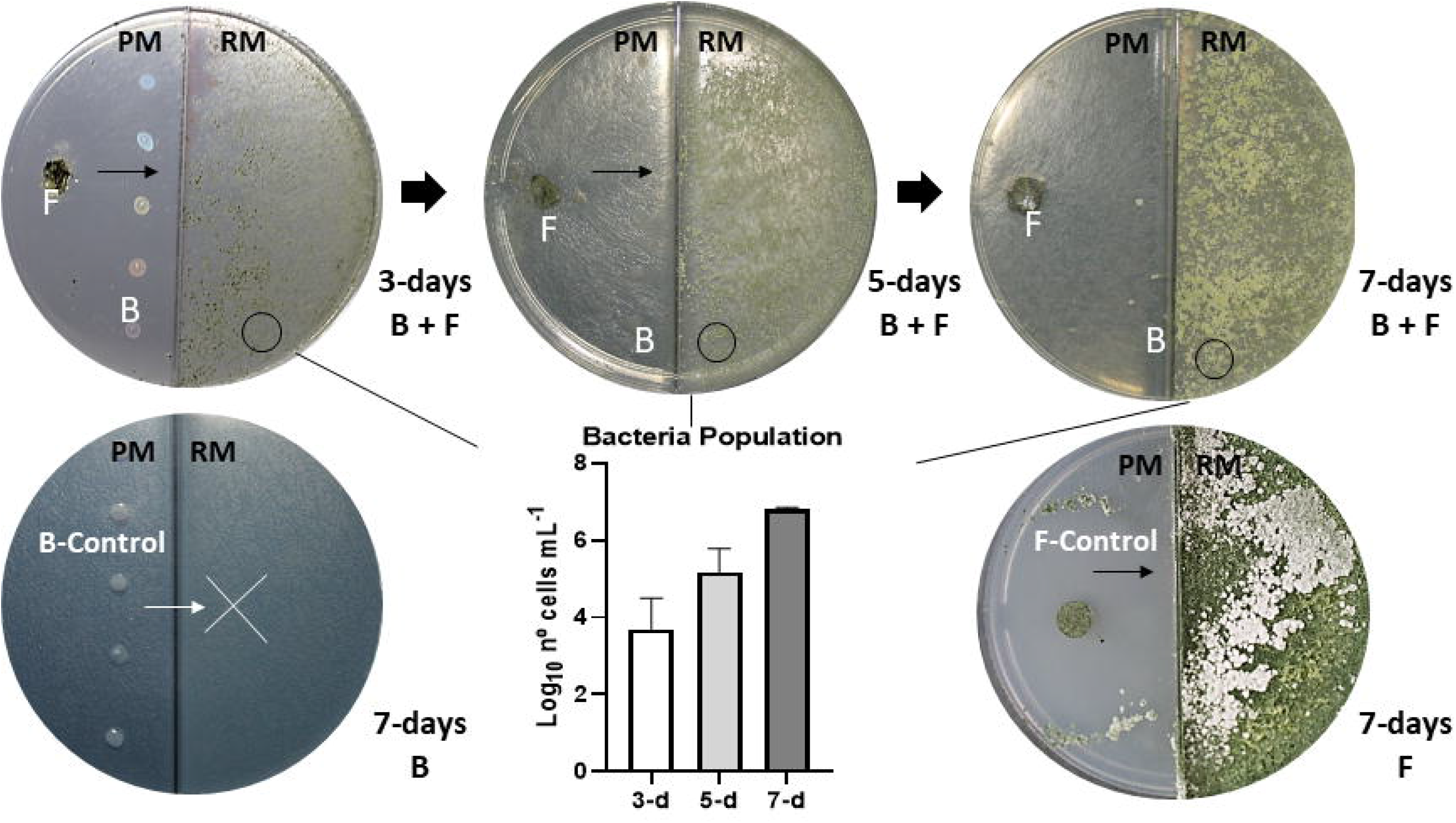
Migration assay in bi-partite Petri dishes with agar plug of fungal hyphae (F) and bacterial suspension spots (B) arranged at the same side of the plate filled with poor nutrient medium (PM). Temporal growth (7-days assay) of the fungus that visible cross the barrier and colonize the rich nutrient medium (RM) compartment. *S. marcescens* population detection at 3, 5, and 7-d at the opposite RM-compartment (expressed in log_10_ CFU. mL^−1^). No *S. marcescens* detection was possible for B-control and F-control at 7-d in the RP-compartment.

### Survival of the *S. marcescens* applied to vermicompost in the presence and absence of the *T. longibrachiatum*

The population density of the *S. marcescens* UENF-22GI on vermicompost in the presence of *T. longibrachiatum* UENF-F476 was accessed over 112-days experimental time (Fig. 6). From an introduced population level of ~10^8^ cells. g^−1^, it was noticed a sharp decline and positive recovery of the bacteria population until the end of the assay (112-d) for both treatments. However, bacteria survival rates were modulated by the fungal presence in the substrate, indicating its influence on the survival of the bacteria. There were no differences in population-level at 14 and 28-d, with survival about 80% related to the initial population size (considered 100%). There was a significant decline of the bacteria population at 56-d when the fungus was absent in the substrate. At 112-d, it was noticed that the bacteria population size was significantly more abundant in the fungal-inoculated substrate related to the control (without fungus). Also, it was not possible to detect *S. marcescens* in non-inoculated substrate over the experimental period (data not shown).

**Fig 6.**
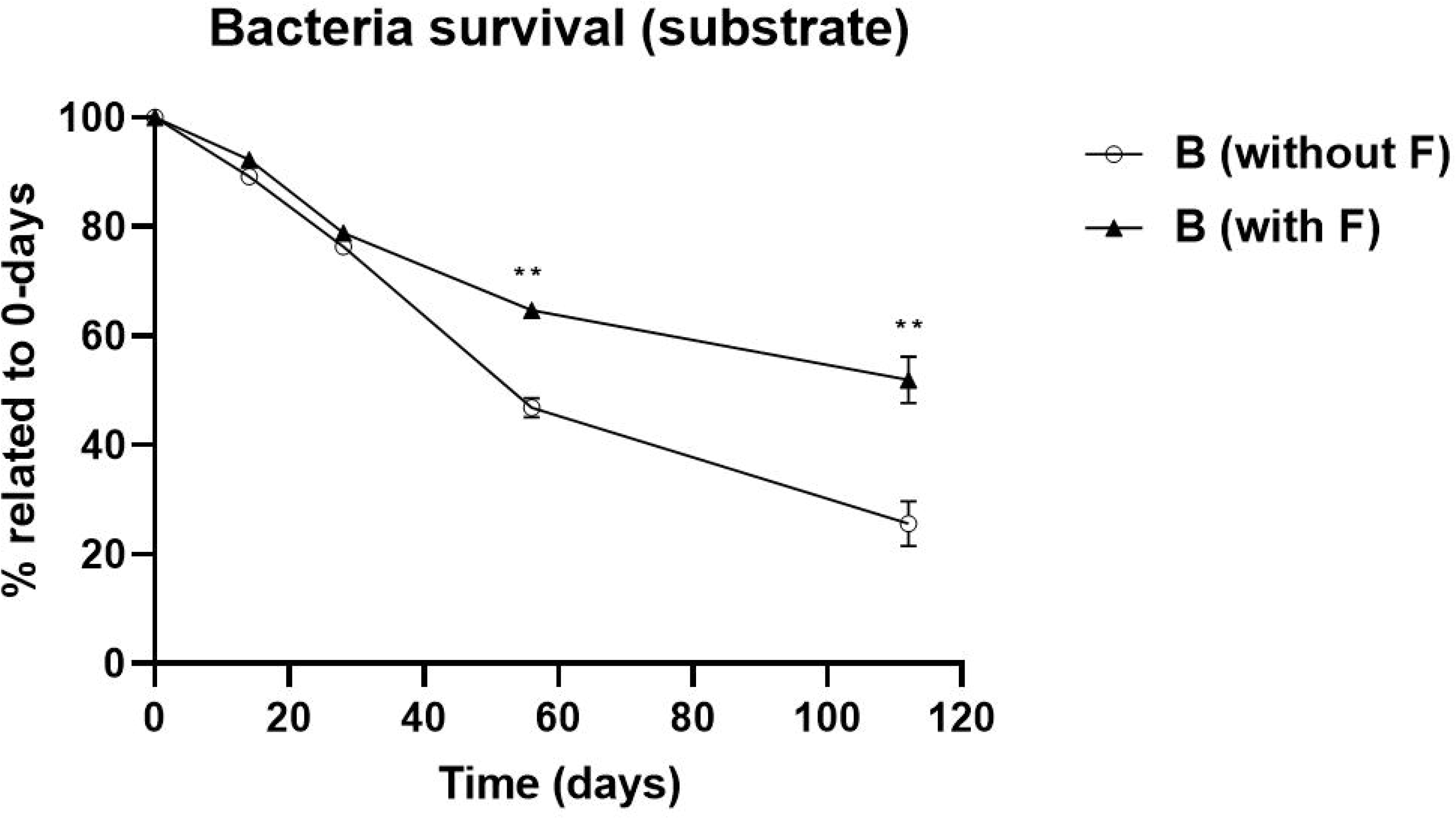
Bacteria (*Serratia marcescens*) survival rates on cattle manure mature vermicompost in the presence or absence of *T. longibrachiatum* fungal co-inoculation over 112-d assay time. The harvest time for bacteria counts was (14, 28, 56, and 112 days after microbial inoculation), and data were expressed as a remaining percentage of the initial bacteria population size (~10^8^ cells g^−1^ substrate equal 100% population level). The pair-wise significance (** p 1%) was obtaining by comparison between B versus B+ F for each harvest time.

### Evaluation of *S. marcescens* plant growth-promoting traits in the presence of the *T. longibrachiatum*

The production of indole compounds was evaluated in the presence of tryptophan as a metabolic precursor of the IAA metabolic route. It was observed that both microorganisms were able to produce indole compounds (Fig.7). When co-cultivated in the liquid medium, there was a significant 40% increase of indole compounds (p ≤ 5%). For P-solubilization, all microbial treatments were able to solubilize the P-Ca source with a range from 149.4 to 851.8 % increase significantly superior to the non-inoculated control. The bacterium *S. marcescens* was superior to *T. longibrachiatum* fungus with ~215 % solubilization activity. The bacteria-fungal combination ability to solubilize P-Ca was synergic increased with 382 and 600% higher than single bacteria and fungus, respectively (Fig. 7). For P-rock solubilization, the bacteria treatment (421.2%) and its combination with the fungus (791.4%) was significantly superior to control and fungus alone. Bacterium-fungi combination was 370.2% superior to bacteria treatment with an apparent synergic effect when both microorganisms were combined (Fig. 7).

**Fig 7.**
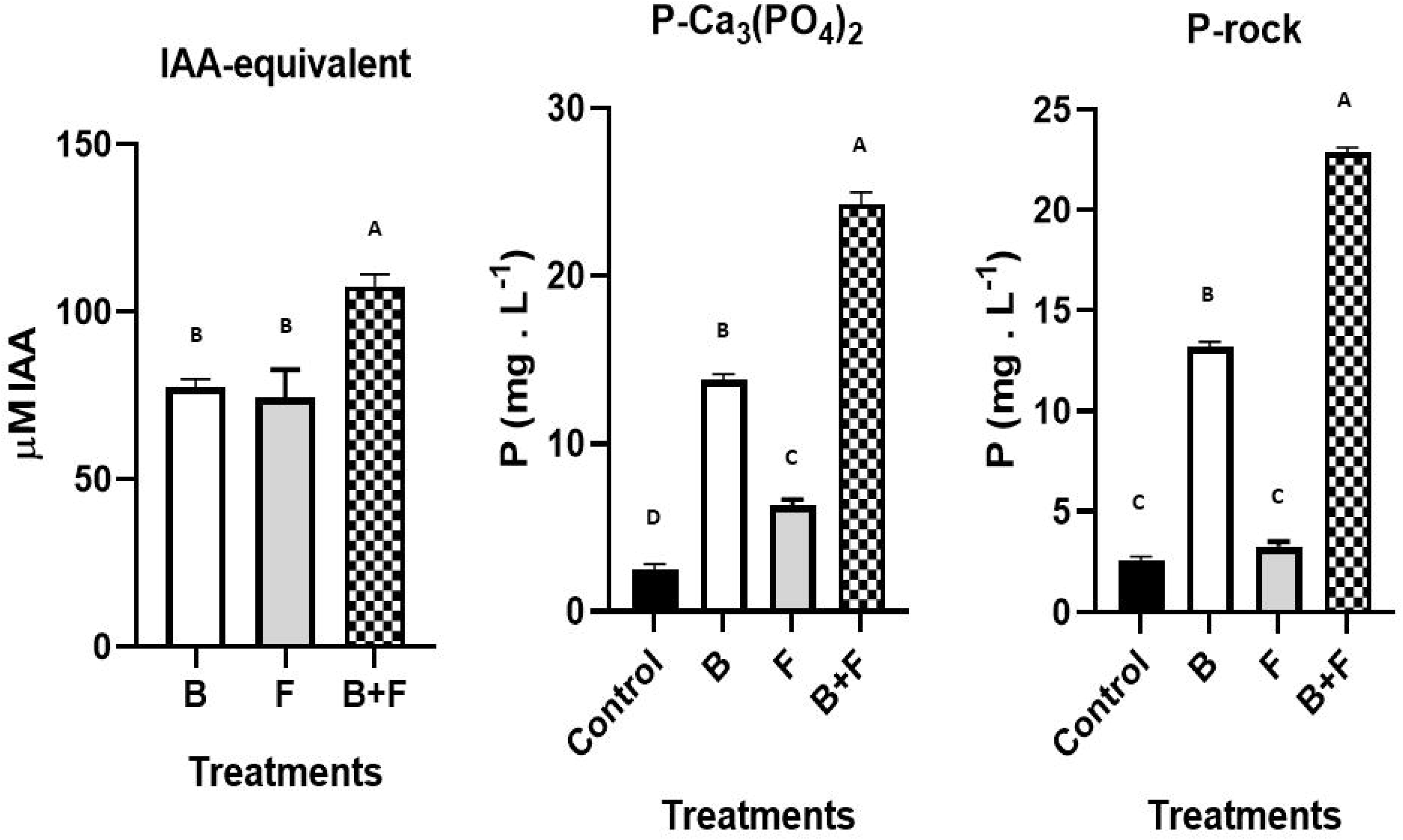
Plant growth promotion traits of *Serratia marcescens* strain UENF-22GI (B), *Trichoderma longibrachiatum* strain UENF-F476 and its combination (B + F) for *in vitro* evaluation of IAA-equivalent production, Ca_3_(PO_4_)_2_ and Araxá P-rock for P-solubilization. Control = non-inoculated. Data represent means of 4 replicates and statistical analysis performed using one-way ANOVA and Tukey test at P < 0.05.

### Plant growth response of the *S. marcescens* substrate-inoculation in the presence of *T. longibrachiatum* under nursery conditions

We accessed the plant-growth response of tomato (Fig. 8) and papaya seedlings (Fig. 9) when vermicompost plant substrate was inoculated with *T. longibrachiatum* UENF-F476 (F), *S. marcescens* UENF-22GI (B), its co-inoculation (B + F) and non-inoculated control (C). For the fresh and dry matter of tomato seedlings roots and shoots, *S. marcescens* inoculation was significantly superior to fungal and control treatments (Fig. 8). A more positive response was observed when the bacterium was applied with *T. longibrachiatum* with increased values ranging from 15 to 60% related to bacteria treatment (Fig. 8). Almost the same trend was obtained when testing papaya plantlets in the same condition. (Fig. 9). Bacteria inoculation was significantly superior to fungus and control treatment for shoot dry matter and root fresh matter. However, the bacteria response combined with the fungal (B + F) had shown positive plant response for all parameters evaluates, with increased values ranging from 15 to 40% related to the bacteria treatment.

**Fig 8.**
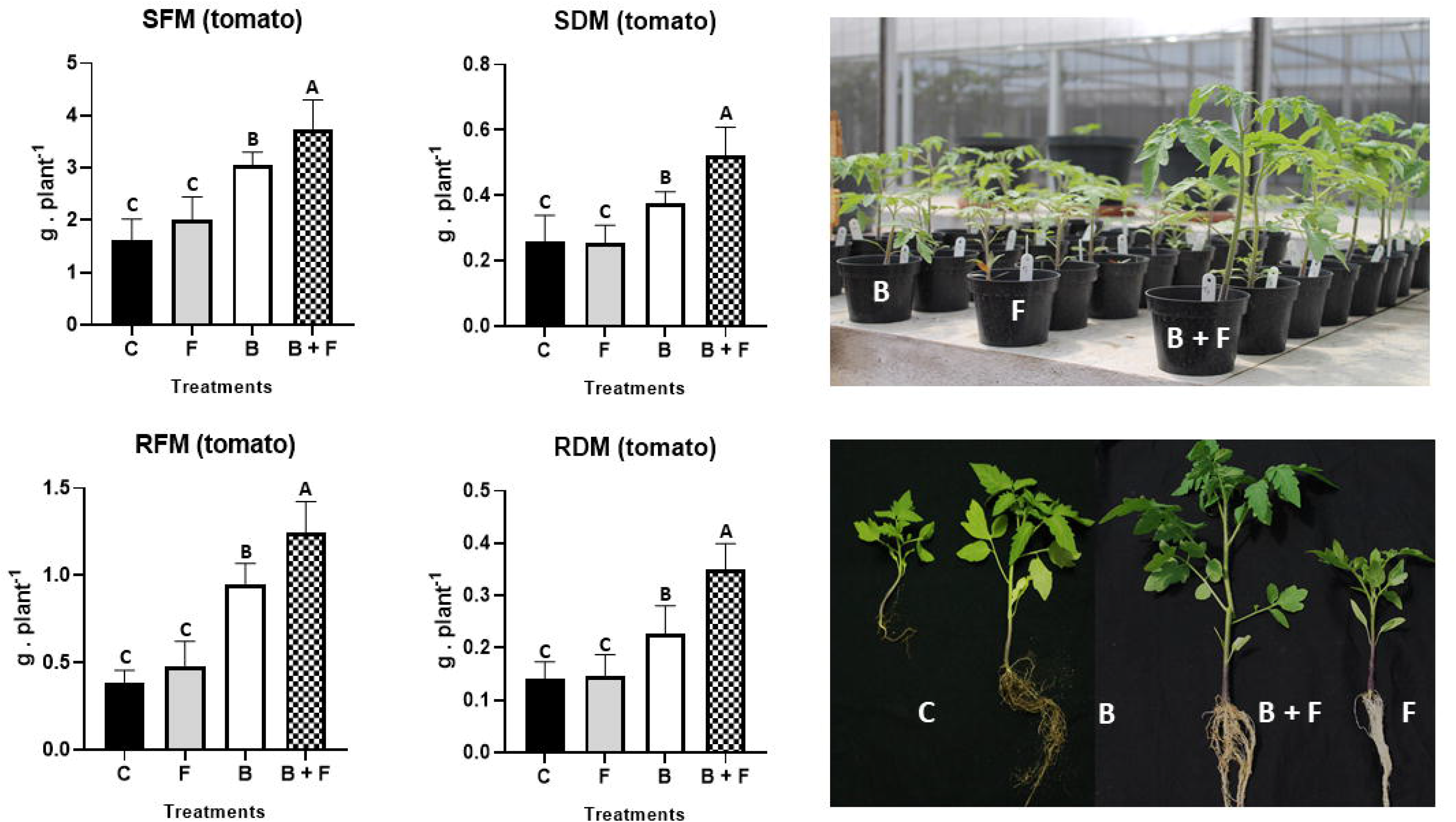
Plant growth promotion response of tomato seedling in substrate enriched with microorganisms. (C) non-inoculated; (F) isolated *Trichoderma longibrachiatum* strain UENF-F476; (B) *Serratia marcescens* strain UENF-22GI, and its combination (B + F) applied to plant substrate containing mature vermicompost. Biometric parameters were shoot fresh matter (SFM), shoot dry matter (SDM), root fresh matter (RFM), and root dry matter (RDM). Graphical Bars with standard deviation followed by the same letter do not differ significantly (p <0.05) by Tukey test.

**Fig 9.**
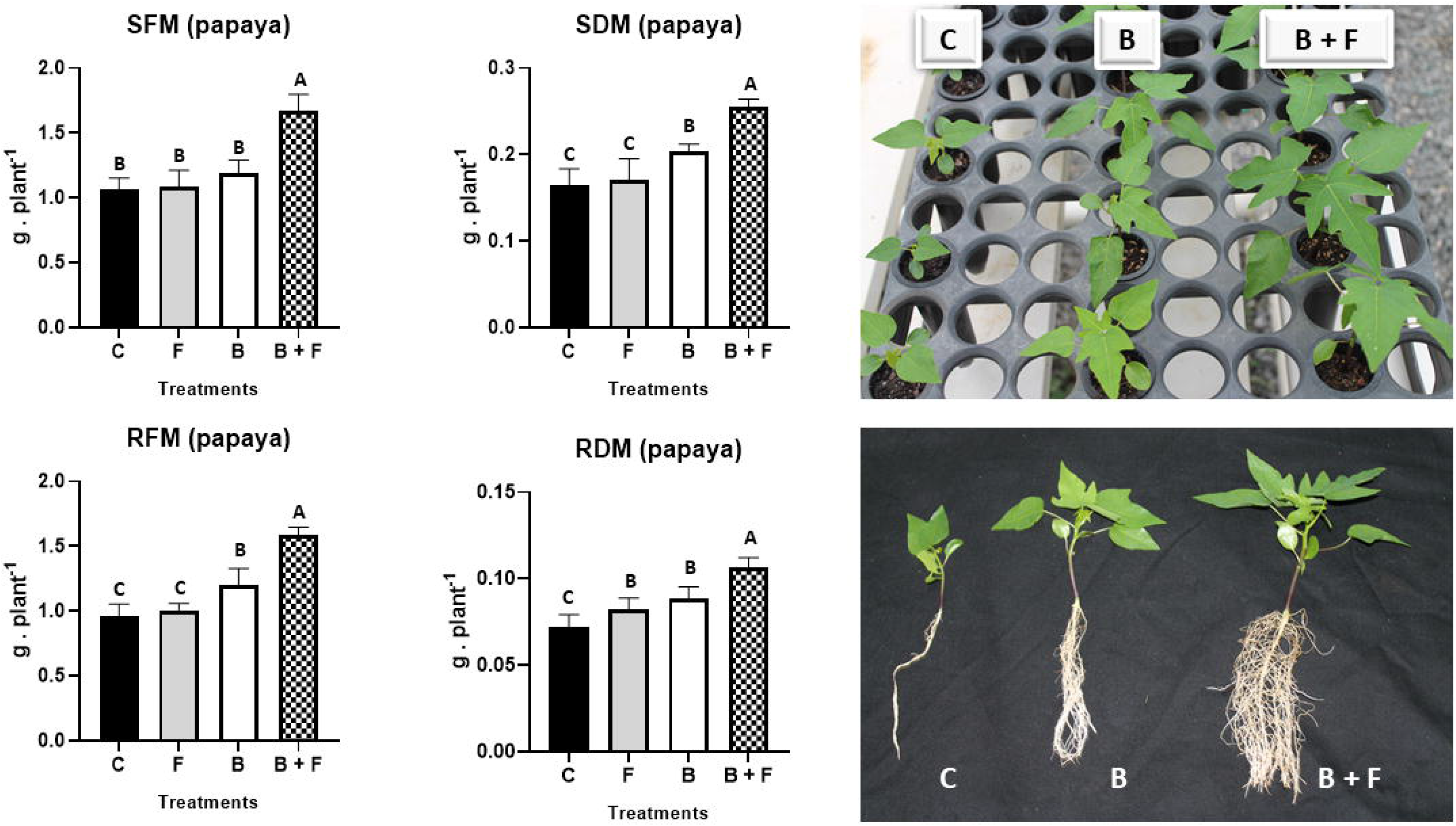
Plant growth promotion response of papaya seedling in substrate enriched with microorganisms. (C) non-inoculated; (F) isolated *Trichoderma longibrachiatum* strain UENF-F476; (B) *Serratia marcescens* strain UENF-22GI, and its combination (B + F) applied to plant substrate containing mature vermicompost. Biometric parameter determined were shoot fresh matter (SFM), shoot dry matter (SDM), root fresh matter (RFM), and root dry matter (RDM). Graphical Bars with standard deviation followed by the same letter do not differ significantly (p <0.05) by Tukey test.

## DISCUSSION

Microbial processes are held by complex communities of prokaryotic and eukaryotic microorganisms in the biosphere. Essential soil-micro-ecosystem services are outcomes from coexisted populations of bacteria and fungi, acting in the biogeochemical cycles, organic matter dynamics, nutrient cycling and soil aggregation (42, 43). Studies involving fungi and bacteria in the soil-systems evidenced a broad range of ecological interactions result in antagonism, competition and mutualistic strategies (44). Pioneer studies emphasizing mycorrhizal fungi interaction with bacteria (14) gave place to a broader scope of mutualistic interactions between saprotrophic fungi and bacteria (12, 21, 23, 26, 45), open up opportunities for a new set of innovative strategies to develop bioproducts for agriculture.

This study hypothesized that a well-characterized beneficial bacterium *Serratia marcescens* UENF-22GI (27) increased its ecological fitness and can improve plant-growth promotion response when combined with a suitable compatible saprotrophic fungus.

The fungal selected (*Trichoderma* sp. isolate UENF-F476)was identified in the present study by phenotypic and molecular approaches as *Trichoderma longibrachiatum*. The *Trichoderma* genus is well-adapted to the soil environment and widely recognized as a bio-control agent (46). The assigned species *T. longibrachiatum* (Fig. 1–2) were also reported as biostimulant agent (47) and cell-wall xylanase producer (48).

We proposed a simple microbiological protocol for screening of suitable microbial-partner that combines colony-colony (Fig. 3) and cell-cell (Fig. 4) interaction analysis. The petri dish filled with NB-solid medium represents a homogeneous rich-nutrient habitat with controlled physic-chemical conditions, where the growth rates and niche occupancy can be easily screened for a variety of microbial combinations. Changes in media composition can achieve a vast array of screening for proper bacterium-fungus combination, abiotic condition, microbial population abundance, and selective pressure (for example, poor-nutrient, specific nutrient deficiency, acidic pH, osmotic pressure, heavy metal contents, etc.). Moreover, the use of media that mimic a target ecological niche (for example, sugarcane apoplast, soil, and compost or vermicompost extract, etc.) represent another exciting approach for target-selection.

The *T. longibrachiatum* and *S. marcescens* colonies were fully compatible, with no sign of antibiosis or niche occupancy restriction (Fig. 3). Also, the species-species specificity was demonstrated when the strain of *S. marcescens* restricted the movement of e phytopathogenic fungus (Fig. 3). *Serratia* genus has been widely recognized for antifungal action related to the chitinolytic activity (49). The genome of Sm strain UENF-22GI harbor four chitinases and other chitin metabolism genes, and the growth-reduction of the phytopathogenic fungi was demonstrated (27). Mutants of *S. marcescens* lacking chitinase expression were also able to kill fungal hyphae, suggesting that other biocontrol mechanisms can operate (50). Matteoli and colleagues (27) proposed that UENF-22GI chitinase activity may interact synergistically with prodigiosin red-pigment and other molecules to suppress fungal growth.

Prodigiosin is the remarkable red pigment of *S. marcescens* UENF-22GI encoded for 14 genes organized in the *pig* operon (27). We observed differential fungal responses to prodigiosin production, where healthy *T. longibrachiatum* hyphae were reddish stained concerning the growth limitation of some phytopathogenic fungi [herein and (27)]. The protective role for the prodigiosin associated with the *T. longibrachiatum* hyphae cannot be ruled out, which could be considered an advantage for fungal competitiveness.

Microscopy analysis of the cell-cell interactions confirmed the microbial compatibility and evidenced the role of the fungal hyphosphere as a microhabitat that harbor bacteria aggregates and biofilms (Fig. 4). Therefore, *S. marcescens* population density and activity was increased by the presence of fungal growth-structures. Also, it was noted that the hyphae tip growth carried small aggregates, promoted migration and dispersion of the bacteria in the environment (Fig. 4–5). The coexistence and mutualistic interaction between bacteria and fungi and their ability to disperse together with hyphae have been wide demonstrated (12, 21, 25, 44).

Nazir and colleagues (45) reviewed the mechanism that promotes the ability of certain bacteria groups to colonize and benefit from the mycosphere. The authors pointed out the efficiency in acquiring specific released nutrients, the presence of type III secretion systems machinery, flagellar movement, and biofilm formation as pivotal aspects of bacterial life in the mycosphere. Biofilms and bacteria dispersion along the hyphae were previously described by (22) that evidenced the ability of the bacterial community to migrate in the soil microcosms through hyphae of the saprophytic fungus *Lyophyllum*. These authors also showed a correlation between migration proficiency and presence of the *hrcR* gene related to the type III secretion system that is involved in bacterial attachment and acted as molecules delivering system to the fungal host.

We observed that the *S. marcescens* was able to recognize, adhere and migrate with *T. longibrachiatum* hyphae, to benefit from C-sources and efficient environmental dispersal. The same *S. marcescens* strain was able to colonize the hyphosphere and migrate on the hyphae of different phytopathogenic fungi, but in this case, limiting their growth. Interestingly, a similar pattern of cell-cell interaction resulted in a different phenotype for *Serratia marcescens*-*Trichoderma longibrachiatum*, where the bacterium does not compromise fungal growth.

The ecological interaction between bacteria-fungi in the soil ranging from antagonistic and mutualistic depending on the microbial species combination (24). We showed that the bacterium-fungi combination evaluated mutually improved its ecological fitness by exchanging benefits and reduce the competition with soil-borne microorganisms. Such ecological concepts can be converted to bioproducts for sustainable agriculture.

The hyphosphere effect can explain the increased survival of S. marcescens co-inoculated with T. longibrachiatum in vermicompost substrate (Fig. 6) overpopulation size and the formation of biofilm-type structures. The biofilm has been considered a prominent survival strategy for successful bacteria colonizers, enabling proliferation in desirable niches by increasing the resistance to biotic and abiotic stresses (51). Structured biofilms can also provide nutritional benefits for members of the community since the extracellular polymeric matrix formed harbor active lytic that enhance nutritional fitness by capture, store and used in times of environmental scarcity (52).

Better survival and dispersion of selected bacteria is one of the critical aspects of the success of the microbial inoculation technology. Avoiding the rapid decline of the bacteria population introduced in seeds, soils, or plant-substrate would increase its beneficial potential for plant growth and development (1). Even, soil-borne bacteria such as the rhizobia, widely used as bioinoculant, had shown improved soil survival when co-inoculated with fungi by forming biofilms (23). Our study demonstrated that *S. marcescens* formed biofilm around the older fungal hyphae and migrate as small aggregates in tip-growing hyphae the colonize new micro-habitats, promoting respectively, survival and dispersion of the bacteria inoculum.

Besides the improvement of bacterial fitness, we also evidenced increase biostimulation and solubilization activity modulated by the fungal (Fig. 7). The ability to solubilize P-sources and produce indole compounds was previously demonstrated (27). The *S. marcescens* UENF-22GI genome analysis evidenced several genes related to the production and secretion of low molecular weight acids that are compounds involved in mineral P-solubilization. Genes related to the IAA biosynthesis pathway were also reported for *S. marcescens* UENF-22GI (27). The plausible hypothesis for the increased activity in co-culture media is the improvement of the bacteria growth (53). We also cannot rule out the altered bacteria-fungal exudates pattern in which a higher amount or more efficient organic acids are released in the culture media, increasing the P-solubilizing activity. Nazir et al. (54) found that the fungus *Lyophyllum* sp., when co-inoculated with *Burkholderia terrae* BS001, exudated higher amounts of compounds. The *in vitro* compatibility evaluated by a simple protocol and increased ecological fitness of the bacterium *S. marcescens* UENF-22GI combined with *Trichoderma longibrachiatum* UENF-F476 serve as a technological platform to develop new bioproducts applied to agricultural systems. The global market for bioproducts is in continuous expansion, and the microbial-based product is the “risen stars” of the market with predictable values of over 10 billion US dollars by 2025 (55). Thus, the technological challenges related to plant response improvement to bacteria bioinoculants can be circumvented co-application with compatible fungus. We analyzed the feasibility of the microbial-partner to promote tomato and papaya growth under nursery conditions. The *Serratia* strains were used as able to promote plant growth in maize plantlets (27). However, when combined with *T. longibrachiatum*, boosted the root, and shoot biomass for both plant species concerning the treatments that received only the bacteria. Here, we delivered the bioproduct for the plants exploring the concept of biological enrichment of plant-substrates (biofortification of plant substrates), but there are open possibilities for formulation and delivery under green-house and open filed conditions.

Mutualistic interaction between mycosphere-colonizing bacterium *S. marcescens* UENF-22GI and the saprotrophic fungi *T. longibrachiatum* UENF-F467 was evaluated by a simple protocol that accessed the biological compatibility between the partners. Such compatibility increased the ecological fitness (substrate survival) of the bacteria alongside with beneficial potential for plant growth. A proper combination and delivery of compatible beneficial bacteria-fungus represent an open avenue for biological enrichment of plant substrates technologies in agricultural systems.

## ACKNOWLEDGMENTS

The authors acknowledge the financial support provided by FAPERJ grant n° E-26/203.003/2017, CNPq grant n° 314263/2018-7, Newton Fund grant BB/N013476/1 “Understanding and Exploiting Biological Nitrogen Fixation for Improvement of Brazilian Agriculture,” co-funded by the Biotechnology and Biological Sciences Research Council (BBSRC) and the Conselho Nacional das Fundações Estaduais de Amparo à Pesquisa (CONFAP) and FINEP-PLURICANA financially supported this study. Also, we are grateful for the strong support given by Dr.Vicente Mussi Dias (LEF/UENF) with the morphological taxonomy of the fungal and Dr. Roselaine Sanchez da Silva de Oliveira that isolate the fungi. This study is part of the Ph.D. of the first author (RJAR), who is grateful for the fellowship conceded by CAPES.

---

**Supplementary Table 01:**
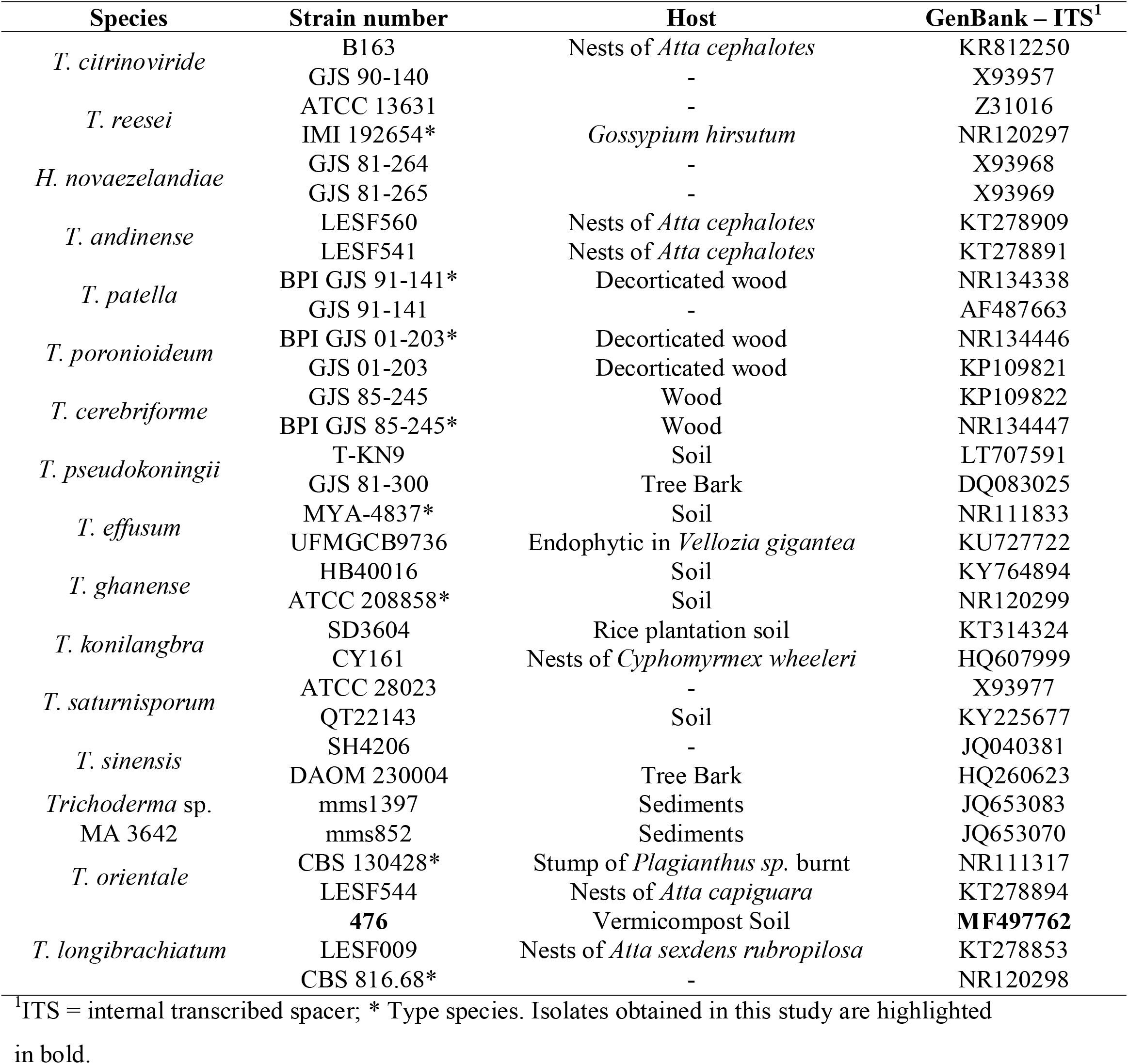
Isolates included in the phylogenetic study of endophytic fungi

